# Experimental Zika Virus Infection in the Pregnant Common Marmoset Induces Spontaneous Fetal Loss and Neurodevelopmental Abnormalities

**DOI:** 10.1101/259317

**Authors:** Maxim Seferovic, Claudia Sánchez-San Martín, Suzette D. Tardif, Julienne Rutherford, Eumenia C.C. Castro, Tony Li, Vida L. Hodara, Laura M. Parodi, Luis Giavedoni, Donna Layne-Colon, Manasi Tamhankar, Shigeo Yagi, Calla Martyn, Kevin Reyes, Melissa Suter, Kjersti M. Aagaard, Charles Y. Chiu, Jean L. Patterson

## Abstract

During its most recent outbreak across the Americas, Zika virus (ZIKV) was surprisingly shown to cause fetal loss and congenital malformations in acutely and chronically infected pregnant women. However, understanding the underlying pathogenesis of ZIKV congenital disease has been hampered by a lack of relevant *in vivo* experimental models. Here we present a candidate New World monkey model of ZIKV infection in pregnant marmosets that faithfully recapitulates human disease. ZIKV inoculation at the human-equivalent of early gestation caused an asymptomatic seroconversion, induction of type I/II interferon-associated genes and proinflammatory cytokines, and persistent viremia and viruria. Spontaneous pregnancy loss was observed 16-18 days post-infection, with extensive active placental viral replication and fetal neurocellular disorganization similar to that seen in humans. These findings underscore the key role of the placenta as a conduit for fetal infection, and demonstrate the utility of marmosets as a highly relevant model for studying congenital ZIKV disease and pregnancy loss.

## INTRODUCTION

Zika virus (ZIKV) is a mosquito-borne (*Aedes* genus) arbovirus of the *Flaviviridae* family. Originally discovered in Uganda in 1947^1^, previous outbreaks of ZIKV were largely sporadic across Southeast Asia and the equatorial African belts, but later spread east resulting in an outbreak in Yap Island in 2007, followed by epidemics in French Polynesia, New Caledonia, the Cook Islands, and Easter Island in 2013 and 2014^1,2^. The virus emerged in the Caribbean and South America in late 2014, persisting currently across the Western Hemisphere as of 2017 (https://www.cdc.gov/zika/geo/index.html). The virus is geographically spread as a result of human travel from endemic regions, alongside human-to-human transmission via sexual intercourse, with blood transfusions, and via vertical maternal-fetal transmission^2–5^

In contrast to other flaviviruses, ZIKV causes fetal loss and malformations which serve as the phenotypic basis for human congenital Zika syndrome (CZS)^8–11^. Although no other flavivirus is known to cause disseminated fetal neural malformations in humans, worldwide concern for latent viral disease was raised following several case reports demonstrating persistent ZIKV RNA in the amniotic fluid, placenta, and fetal neural tissue weeks to months after initial maternal infection^3,4^. While 80% of pregnancies remain asymptomatic, a minority of patients experience a self-resolving acute illness characterized by fever, rash, and/or conjunctivitis^5^. ZIKV has also been associated with acute neurological illnesses in children and adults, including meningoencephalitis and acute flaccid paralysis, as well as Guillain-Barré syndrome^6^.

A number of animal models of ZIKV infection have been described to date. These include several murine models in which ZIKV-associated complications in pregnant females required the use of immunodeficient animals with defects in the interferon-related signaling pathways^7–10^. Rhesus, pigtail, and cynomolgus macaque models of ZIKV infection have also been developed^11–15^. Acute ZIKV infection in these nonhuman primate (NHP) models early in the course of the pregnancy have been shown to result in an average spontaneous abortion rate of 38% (Dudley, *et al.*, under review), with one study in a single pigtailed macaque demonstrating impaired neurodevelopment associated with ZIKV infection of the fetal brain and neuroprogenitor cells^11^. However, there are several limitations to existing NHP models of viral congenital infections, including long lengths of gestation, limited mating periods associated with long intervals of ovulatory suppression between pregnancies, and a large body size that makes higher biocontainment housing and testing of limited-quantity vaccines or therapeutics challenging.

In contrast, mounting evidence over the past decade has pointed to the common marmoset (*Callithrix jacchus)*, a rat-sized, neotropical primate, as a relevant and convenient model for investigating viral pathogenesis^16^. The common marmoset model has been shown to reproduce the human illnesses caused by Lassa virus^17^, Ebola / Marburg virus^18^, and flavivirus (dengue virus and West Nile virus)^19,20^ infection. We also recently reported that the course of infection in ZIKV-infected male marmosets resembles that seen in acutely infected humans in many aspects^21^. Specifically, viral illness was subclinical, the virus persisted in body fluids such as semen and saliva for longer periods of time than in serum, and there were neutralizing antibody as well as characteristic interferon-associated host responses. Encouraged by these initial studies demonstrating that viral illness in male marmosets is akin to the human condition, we sought to determine whether infection of marmoset dams would more faithfully recapitulate maternal infection and CZS in humans when compared to murine models and provide key opportunities not afforded by other NHP models. Moreover, we hypothesized that crucial mechanistic insights pertaining to placental infection and vertical transmission could be reached as a result of the unusual but well described characteristics of trophoblast and embryonic development in the common marmoset. Here we report the outcomes of experimental infection with ZIKV in two pregnant marmoset dams, and the resultant impact on the placenta and developing fetus.

## RESULTS

### Experimental ZIKV infection of pregnant marmosets

We inoculated 2 pregnant marmosets (dams) with Brazil ZIKV strain SPH2015 (an Asian lineage strain first isolated in 2015) at estimated gestational days 79 (dam 1) and 68 (dam 2) with a dose at 2.5×10^5^ plaque-forming units (PFU), followed by a second inoculation with the same dose 4 days later (**Fig. 1**). Gestational age prior to inoculation was confirmed by fetal sonogram with biometry and crown-rump length (CRL) measurements. The chosen gestational windows span the ongoing and prolonged periods of placentation in marmosets, as well as critical phases of embryonic neurodevelopment (Carnegie embryonic stages 16–23; **Fig. 2**). Both animals were assessed daily for evidence of clinical signs of illness, and remained asymptomatic following inoculation until delivery. Spontaneous expulsion of intrauterine demised fetuses at days 16 and 18 post-inoculation occurred for dams 1 and 2, respectively (**Fig. 1A**). Fetuses (dichorionic, diamniotic twins for dam 1, singleton for dam 2) were documented via ultrasonography to be alive 2 and 4 days prior to fetal demise and expulsion, respectively.

**Figure 1.**
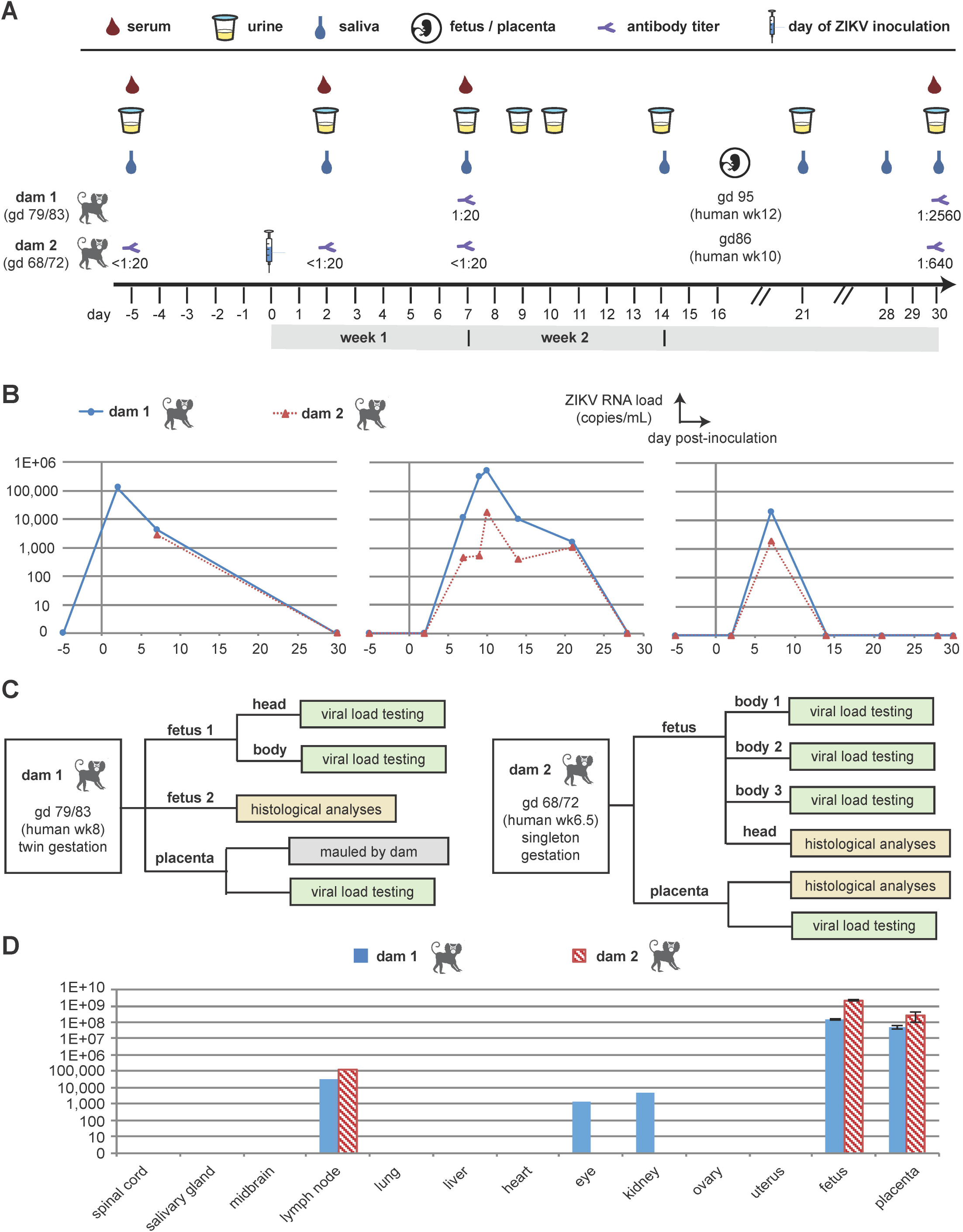
Study overview and results from viral load and antibody testing. (A) Timeline showing days of sample collection before and following ZIKV inoculation at day 0, including collection of placental and fetal samples at time of fetal demise and expulsion (16–18 days post-inoculation). Measured Ab titers at days 7 and 30 post-inoculation for dam 1, and at days −5, 2, 7, and 30 post-inoculation for dam 2 are also given. (B) Viral loads in serially collected body fluids. (C) Flow chart describing analysis of placental and fetal samples. (D) Viral loads in maternal tissues at time of necropsy, and in fetal / placental tissues at time of fetal demise. The error bars shown for the fetal/placental viral loads are based on multiple tissue replicates. Abbreviations: gd, gestational day; wk, week.

**Figure 2.**
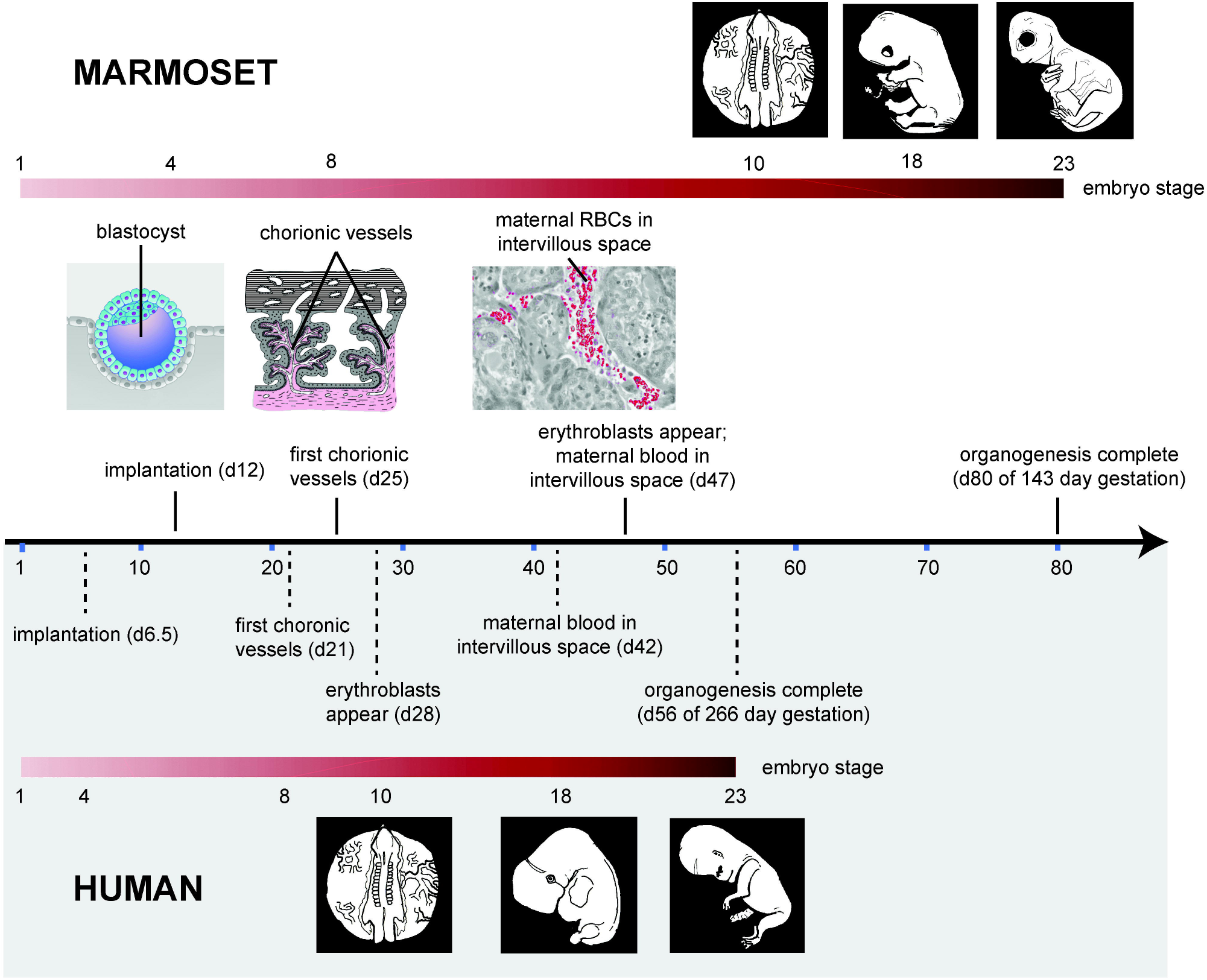
Comparison of the relative pace of placental and embryonic development in humans and marmosets. The embryonic stages shown encompass those that occurred in the Zika-infected marmoset embryos and reflect critical neurodevelopmental stages. The human blastocyst implants on day 6 post-conception, while in the marmoset implantation occurs on day 12. Marmoset organogenesis is complete by approximately day 80^28,50^, compared to day 56 (end of the 8^th^ week) in humans, marking the shift from embryo to fetus. The first appearance of chorionic vessels in the human placenta occurs in week 3^51^; the corresponding period of 20–25 days is when vessels appear in the marmoset placenta^28^. Erythroblasts first appear in human chorionic vessels around day 23, i.e. early week 4, the equivalent to marmoset day 36. In this regard, the marmoset may be delayed, because erythroblasts inside chorionic vessels are not reported to occur until days 46–50. In the human, maternal blood supply is not fully established until week 12, but initial entry of maternal blood into the intervillous space occurs as early as mid-week 6. In the marmoset placenta, maternal blood is first apparent on day 45, equivalent to late week 5 in the human placenta. In New World monkeys such as marmosets, placental interdigitation is described as “trabecular” with cytoplasmic bridges instead of “villous” with finger-like appendages. As this minor anatomical difference is not known to be functionally significant, for clarity of comparison the terms “villous” and “intervillous spaces” are used to describe the marmoset placenta.

### Viral load and serological analyses

Serum, saliva, and urine samples were collected longitudinally at *a priori*-determined specific intervals, including a pre-infection sample taken 5 days before infection, and for up to 30 days following inoculation (**Fig. 1A**). Viral RNA loads were measured in body fluids and tissues by quantitative reverse transcription polymerase chain reaction (qRT-PCR) testing (**Fig. 1A**). Detectable viral loads up to 3 weeks post-inoculation and seroconversion by plaque reduction neutralization testing (PRNT) confirmed both active viral replication and acquired adaptive immunity, respectively Specifically, ZIKV was detectable in the serum for at least 1 week following inoculation by qRT-PCR, with a day 2 peak viral load of ~10^5^ copies/mL, while ZIKV levels in urine and saliva peaked at later time points (day 10 and day 7) and persisted for 2 and 3 weeks respectively (**Fig. 1B**). Both marmosets were seronegative prior to infection (Ab titers <1:20) and seroconverted at 30 days post-infection with positive titers of 1:2560 for dam 1 and 1:640 for dam 2 (**Fig. 1A**).

Maternal necropsy at day 30 post-inoculation demonstrated absence of viral RNA in all tested body fluids, including blood, urine, and saliva. This corresponded to 12 and 10 days post-pregnancy loss for dams 1 and 2, respectively. However, ZIKV RNA was detected by nucleic acid testing (NAT) in maternal lymph node, ocular, and renal tissues, as well as in placental tissues and fetal brain/cranium and abdominal tissues (**Fig. 1C and D**).

### Gross and histological features of placenta and aborted fetuses

Two days prior to fetal demise and expulsion, dam 1 was confirmed by fetal sonogram to carry a live dichorionic, diamniotic twin gestation with normal amniotic fluid volumes and cardiac activity. The spontaneous demise and delivery of the twin fetuses and their placentae were discovered approximately 20-36 hours after expulsion (3 days post sonogram). Both of the aborted fetuses were friable and each had a measured biparietal diameter (BPD) that were 50% of the expected values based on the ultrasounds performed 2 and 4 days earlier; however, this is most likely due to cranial collapse and dolicocephaly typical in fetal tissue after prolonged (>4-6 hours) demise. One twin fetus was immediately placed in formalin for the histological analyses, while the other fetus was dissected for RNA viral load testing (**Fig. 1C**). The placenta was mauled by the dam and unsuitable for histological analysis.

For dam 2, early gestational sonogram at 56 days, prior to ZIKV inoculation, revealed a singleton gestation, an indication of early intrauterine demise of one of the fetuses that occurs in approximately 10-15% of marmoset pregnancies^22^. Thus, although marmosets are obligate carriers of multi-fetal gestations due to a polygenic set of synonymous and non-synonymous substitutions^52,53^, in a number of instances they have been shown to carry singleton fetuses successfully to term^22^. After confirming a continuing developing viable singleton gestation, dam 2 was infected with 2 ZIKV inoculums on days 68 and 72. A viable singleton gestation was confirmed on day 82 and subsequently demised with fetal expulsion on day 86. The single expulsed fetus and its placenta were immediately recovered; the head was formalin-fixed for histology while the body was dissected for RNA viral load testing (**Fig. 1C**). Formalin-fixed paraffin-embedded (FFPE) sections of the placenta alongside positive and negative controls were assessed for the presence of ZIKV using 4G2 antibody against a flavivirus envelope protein (**Fig. 3A & B**). The villous parenchyma stained strongly positive for ZIKV protein by immunohistochemistry (**Fig. 3B**). *In situ* hybridization (ISH), targeting positive-strand ZIKV RNA and its corresponding negative-strand replicative intermediate, established the presence of viral genomic RNA and active replication within placental tissue, respectively (**Fig. 4A & B**). Positive signal for genomic ZIKV RNA strands was adjacent to the nucleus, consistent with the establishment of flaviviral replication factories in the endoplasmic reticulum^23^. The ZIKV protein immuno-staining overlapped with the hybridization positive and negative strand signals, with comparatively much stronger signal from the stable viral genomic strand (+) compared to the replication (-) strand (**Fig. 4B**). Despite clear evidence of extensive placental infection by ZIKV, examination of histological sections stained with hematoxylin and eosin (H&E) revealed no leukocyte infiltration and inflammation in the placenta (**Fig. 4C**); this is consistent with absence of placental inflammation observed in human CZS^50,51^.

**Figure 3.**
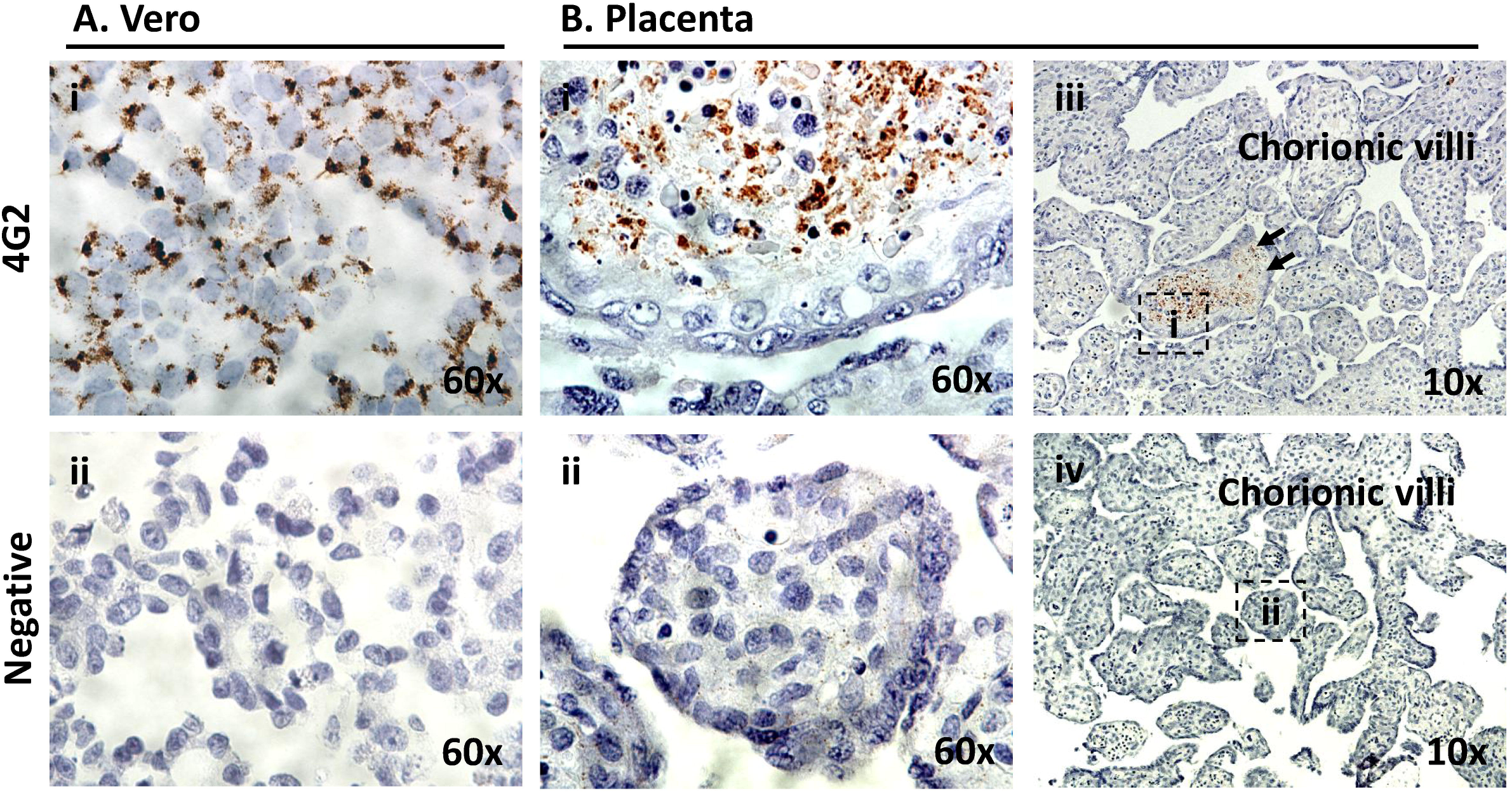
Immunohistochemical 4G2 antibody labeling against Flavivirus E-protein in the placenta of a ZIKV infected marmoset dam. The pregnant adult marmoset (dam 2) was inoculated at days 79 and 83 of gestation (human equivalent of 14 weeks). Placental tissue was collected upon expulsion following spontaneous abortion occurring 16 days post-infection (DPI). (A) Vero cells replicating ZIKV were fixed at 6 DPI as positive control (4G2) (Ai); to control for non-specific labeling, ZIKV infected Vero cells were probed with only secondary antibody (negative) (Aii). (B) 4G2-specific chromogenic signal is apparent with in situ labeling of marmoset placental villi in the parenchyma (Bi) but not negative controls (Bii, iv). Areas imaged at 60x high power (B, i-ii) are indicated by dashed boxes in B iii and iv (4G2 labeled and negative control, respectively).

**Figure 4.**
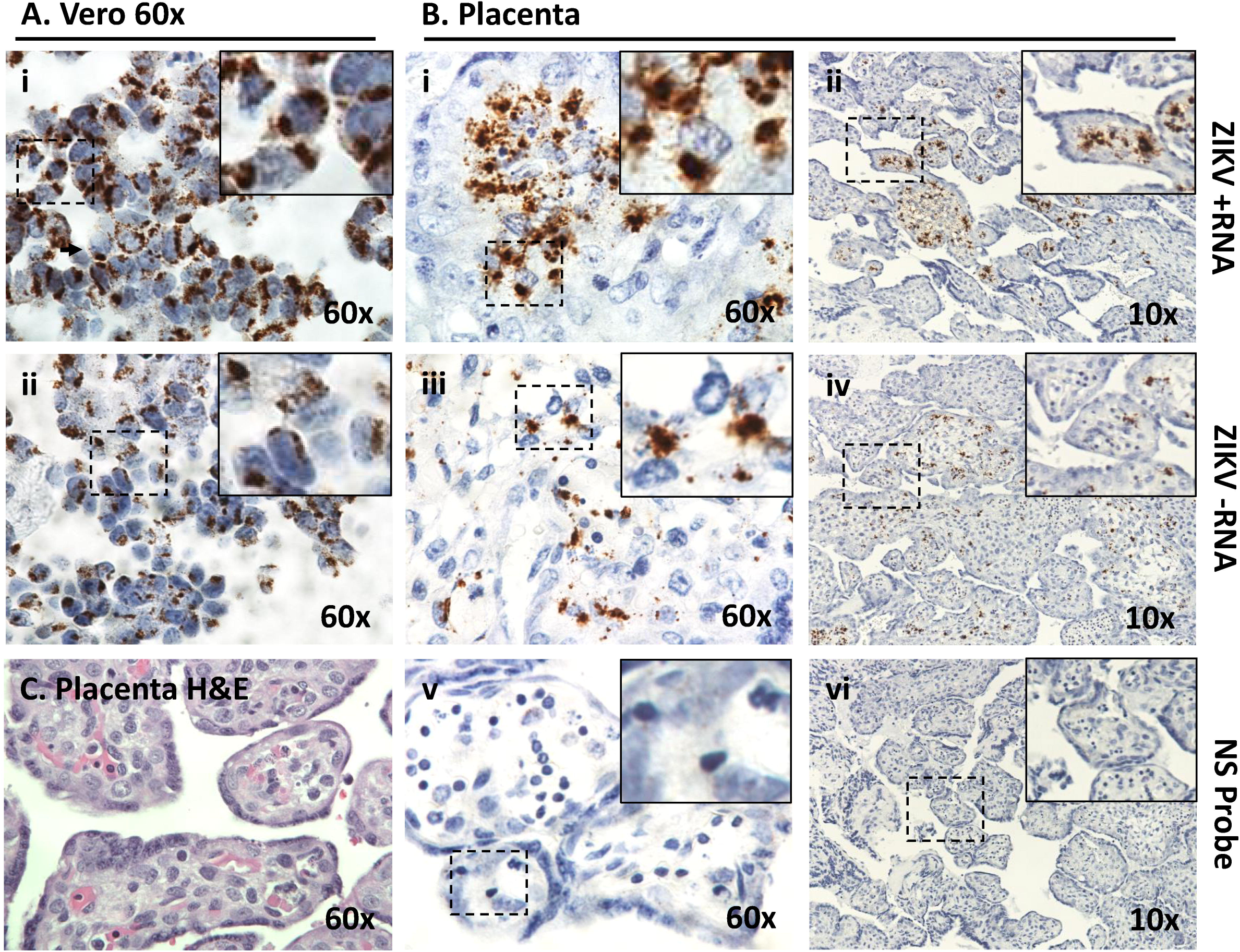
*In situ* hybridization + and – single strand RNA (ssRNA) oligonucleotide probes in spontaneously aborted marmoset (dam 2) placental tissue. The concomitant labeling of both + and – ssRNA is indicative of actively replicating ZIKV. Villous sections were hybridized using amplified probe sets against the +ssRNA the −ssRNA genome to localize active viral replication in the placenta. Fixed Vero cells actively replicating ZIKV at 6 DPI were used as positive controls (A). Placental tissue sections were highly positive for the presence of genomic RNA in villous parenchyma (B, i-ii). Furthermore, ubiquitous detection via ISH of the negative replication strand indicated ZIKV as established in the fetal-placental tissue of the chorionic villi (B, iii-iv). A probe set against human peptidylprolyl isomerase (PPID) mRNA was used as a negative control (B, v-vi). (C) Examination of H&E stained sections of the placental villous revealed no leukocyte infiltration. Dashed boxes denote the zoomed area depicted at high power (60X).

Having established that the placenta in dam 2 actively harbored and propagated ZIKV, we next examined the abdomen and thorax from the dam 1 fetus for evidence of infection using the ISH (+) and (-) strand probes, reserving the corresponding abdomen and thorax from the dam 2 fetus for viral load testing (**Figs. 1C** & **5**). H&E stained sections were used to identify the primitive organs and tissues of the developing fetus (**Fig. 5A**). The hepatic tissue of the animal appeared highly autolyzed, consistent with the prolonged 20-36 hour post-expulsion period, but other tissues and structures were well preserved, with bone or developing cartilage of vertebrae and ribs, and surrounding muscle and primitive mesenchymal tissue apparent and readily identifiable (**Fig. 5Ai**). The fetal thorax and abdomen were diffusely positive for ZIKV hybridization of both + and – RNA strands, with the strongest signal in liver tissue (**Fig. 5Bii**); however, ZIKV-specific hybridization of either strand was not stronger than that measured in the placenta (**Fig. 4B**). Differentiated bone and cartilage did not hybridize with either positive or negative strand probes of ZIKV, whereas viral infection of the surrounding muscle and mesenchymal progenitor tissue was apparent (**Fig. 5B & C**) with evident active viral replication (- strand ISH, **Fig. 5A–C**, **iii**). As with placenta, there was no evidence of inflammation in the fetal tissues despite the apparent extensive infection (**Figs. 4C & 5**).

**Figure 5.**
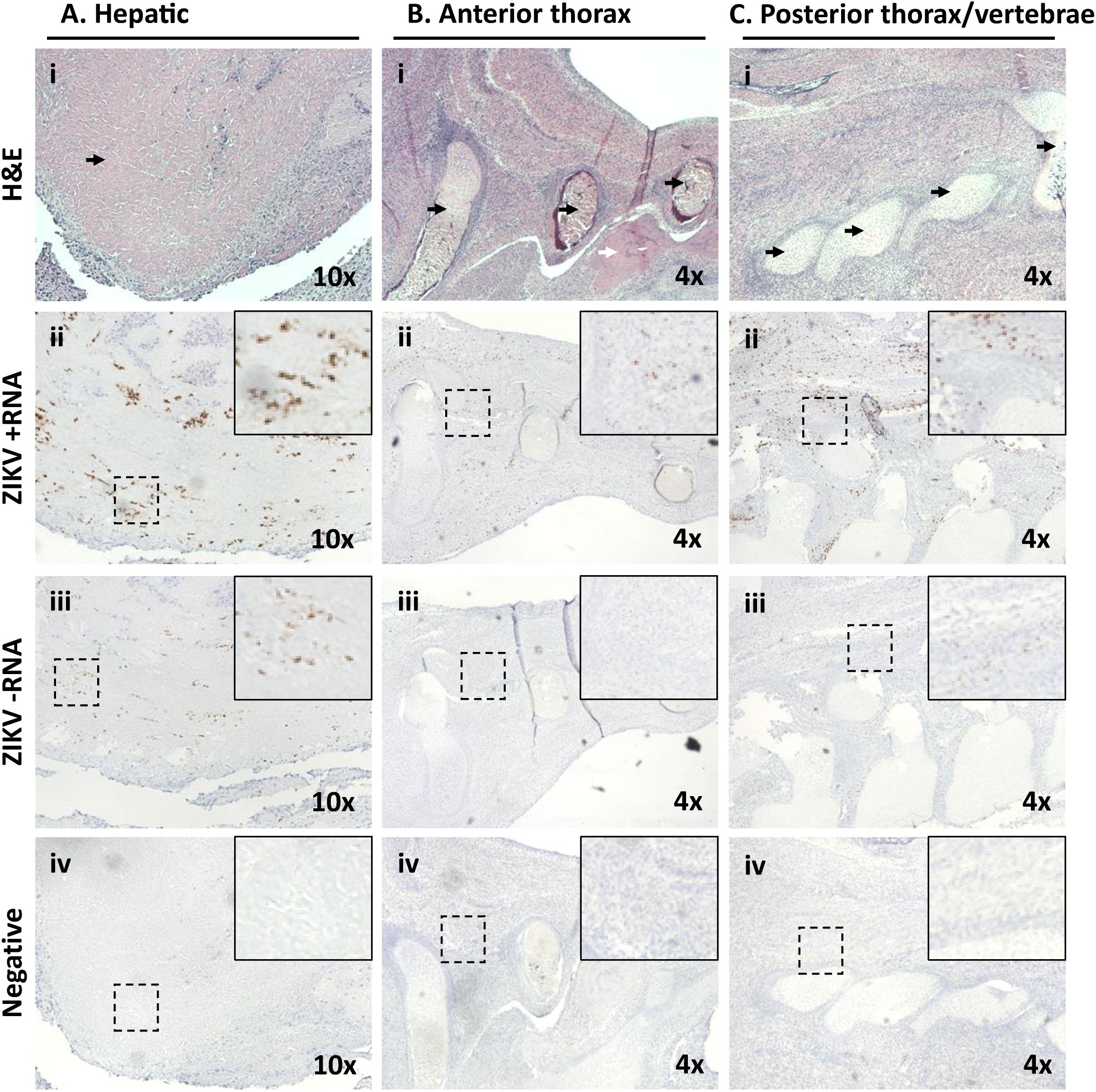
Maternal ZIKV infection during placentation in the marmoset yields later limited active viral replication in the fetal thorax and abdomen, with fetal expulsion 14 DPI. Dam 1 was inoculated at days 68 and 72 of gestation (human equivalent of 14 weeks). Morphometric analysis of H&E stained sections revealed tissue and organ typing, with labeling of ZIKV +ssRNA and −ssRNA localized regions of active viral replication. Overall signal was diffuse across many tissues but was not stronger than placenta (Figure 2). (A) Histological sections of the abdomen demonstrate hepatic autolysis (i, arrow) with strong +ssRNA and −ssRNA ZIKV labeling indicative of active viral replication (ii-iii). (B) Microscopic sections of the superior right hypochondriac anterior thorax (rib cartilage, black arrows) just superior to the liver tissue (white arrow) (Bi) with scant evidence of active viral replication by – ssRNA labeling the primitive mesenchymal and muscle tissue around the rib show signal (B, ii-iii). (C) Sections around the developing cervical vertebrae in the posterior thorax (i, arrows) with rare dual ssRNA labeling indicative of scant active ZIKV replication in the muscle and immature mesenchymal tissue. Hybridization probe-free serial-sections were used as negative controls. Dashed boxes denote the zoomed area depicted.

Given that the most prominent malformations associated with CZS, including microcephaly, are ocular or neurologic^24^, close examination of fetal cranium and its contents from both dam 1 and dam 2 fetuses was made to characterize and anatomically localize cephalic viral infection and replication. The ophthalmic and cerebral regions of a frontally sectioned head of dam 2 fetus were examined (**Fig. 6**). ZIKV signal from both positive viral genomic and negative replication strands was present behind the eye in the muscle and surrounding tissue (**Fig. 6A**, **arrows**). Among the strongest cephalic signal was from the perineurium region of an intra-orbital nerve, likely ophthalmic (**Fig. 6B**). Neuroprogenitor cells (NP) of the developing cortical plate were also highly infected and shown to be actively replicating ZIKV (**Fig. 6C**, **arrows**). Similarly, the optic nerve and anatomy of the eye was most clearly visualized from a sagittal H&E stained section in the fetus of dam 1 (**Fig. 7A**), with the strongest ZIKV-ISH signal in the fetal head occurring in ocular muscle (**Fig. 7B**) and with evidence of active viral replication (**Fig. 7C**). Although the ZIKV signal was weaker in the brain of the fetus from dam 1 relative to the fetus from dam 2, areas of infection and active ZIKV replication were clearly observed (**Figs. 6 & 7**), with H&E sections showing regions of disorganization within the cortical neurons indicating a disruption in development and neuronal migration patterns of the cerebral cortex (**Fig. 8**).

**Figure 6.**
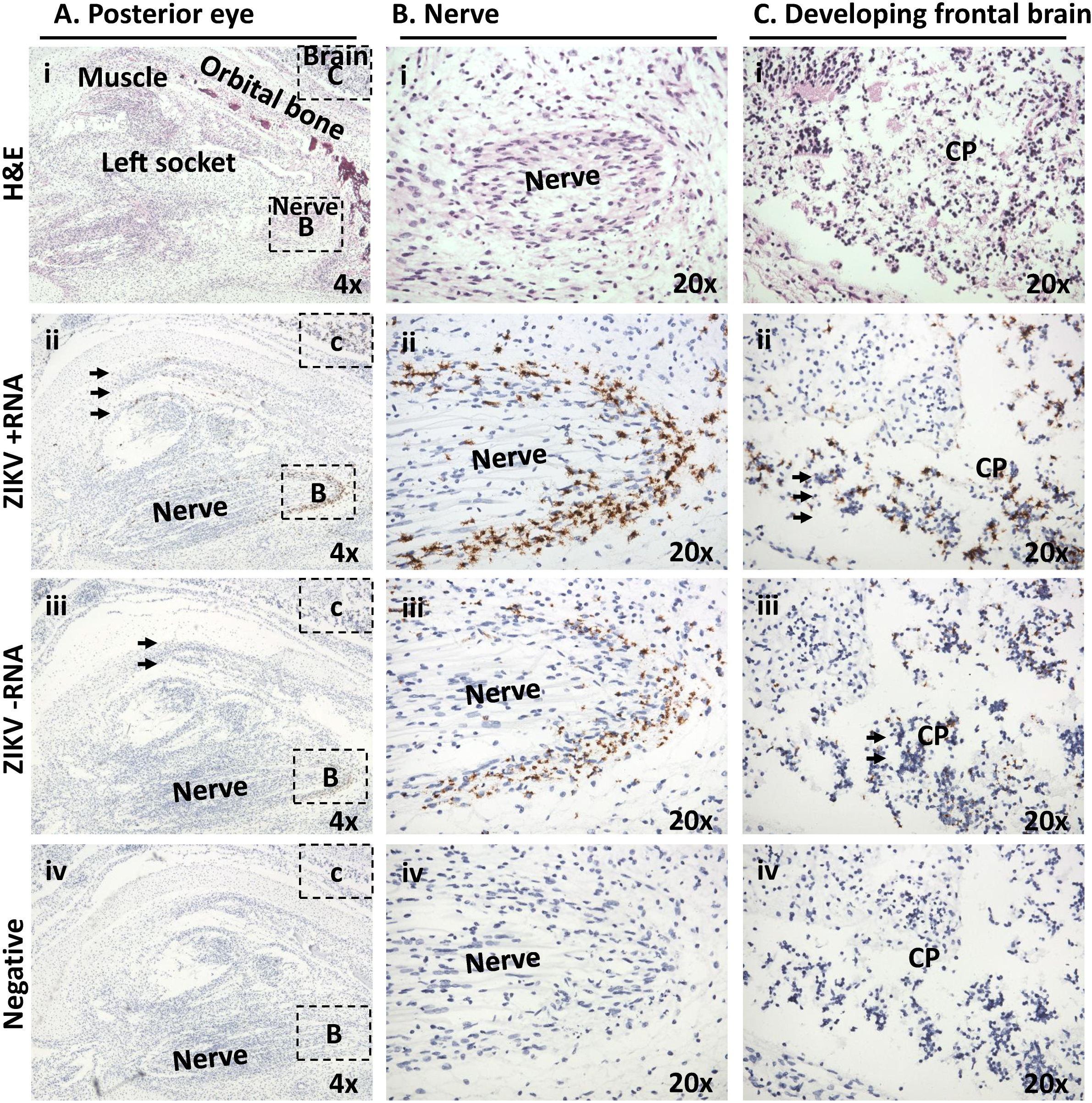
Maternal ZIKV infection during placentation renders significant latter neurophthalmic viral replication. Tissues in the fetus from dam 2 were identified based on morphology using H&E stained sections under low (A) and medium power (B-C). (A) Shown is a frontal cross section taken immediately posterior to the eye. Visible features are identified in an H&E stained section, including orbital bone, ocular muscles and nerves, as well as cortical plate of the frontal brain (i). Areas immediately posterior to the eye, inclusive of the ocular muscle and socket, demonstrated active viral replication as measured by dual labeling of both ssRNA strands (ii-iii, arrows). (B) A boxed area from the same slide at higher magnification showing nerve. Although relatively acellular, the perineurium is constituted of macrophage-like epithelioid myofibroblasts (i, dashed line). Perineural cells around a nerve focally showed the highest replication (ii,iii). (C) A boxed area at higher magnification showing brain. Cortical plate (CP) neurons are seen to be strongly infected (arrows).

**Figure 7.**
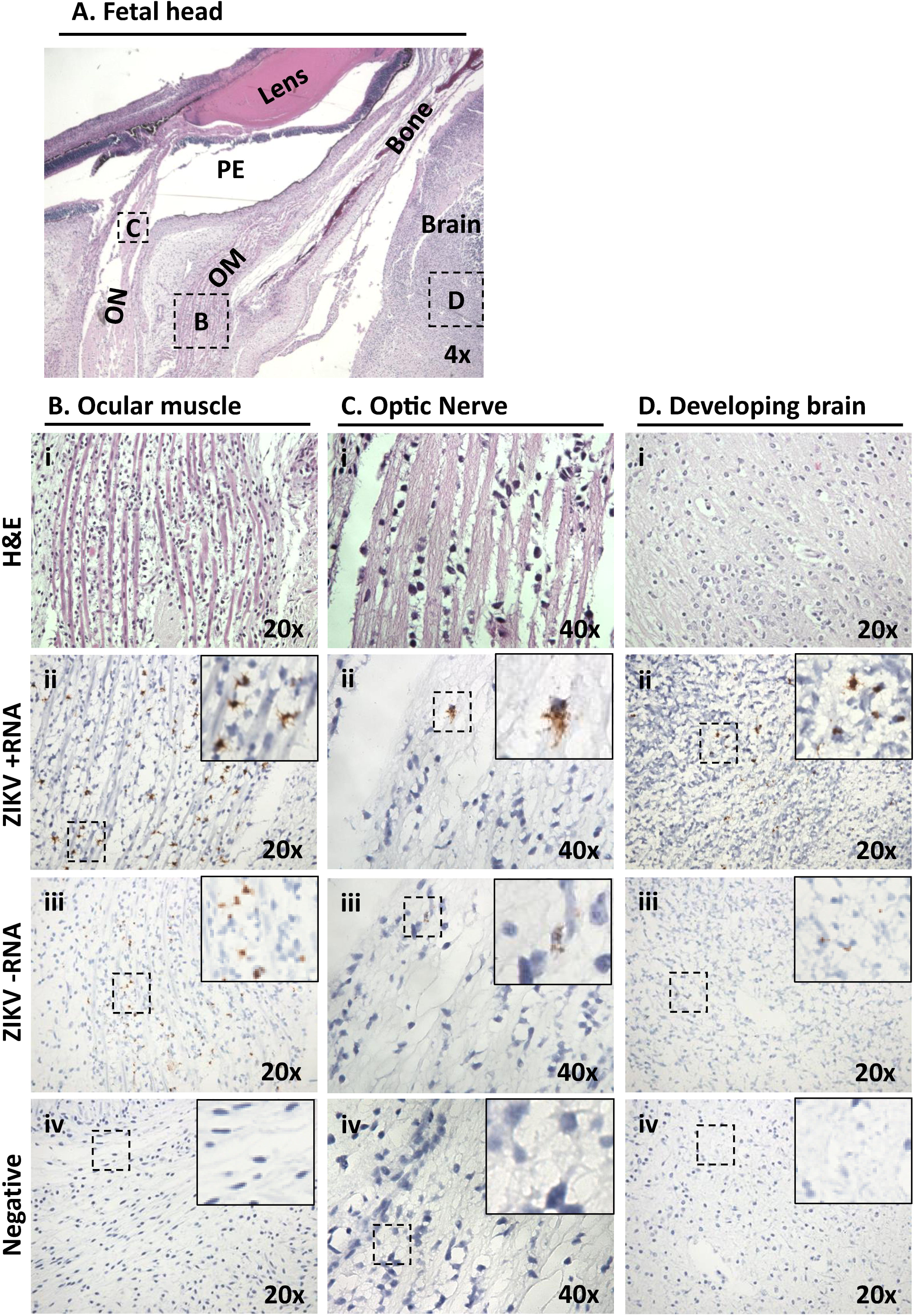
ZIKV infected dam 1 fetus during latter stages of placentation demonstrates ongoing vertical maternal-fetal transmission with active replication in the frontal brain and fetal eye. (A) Low-power H&E stained sagittal cross-section provided anatomic orientation for dam 1 fetus. The primitive eye (PE) with retinal pigment, eye muscle (EM) and the optic nerve (ON) are identified. Boxed areas indicate the location of ZIKV probed in serial sections (B-D). The most highly infected area within the fetal head cross section was found to be the mesenchymal cells of the ocular muscle (Bii,iii).

**Figure 8.**
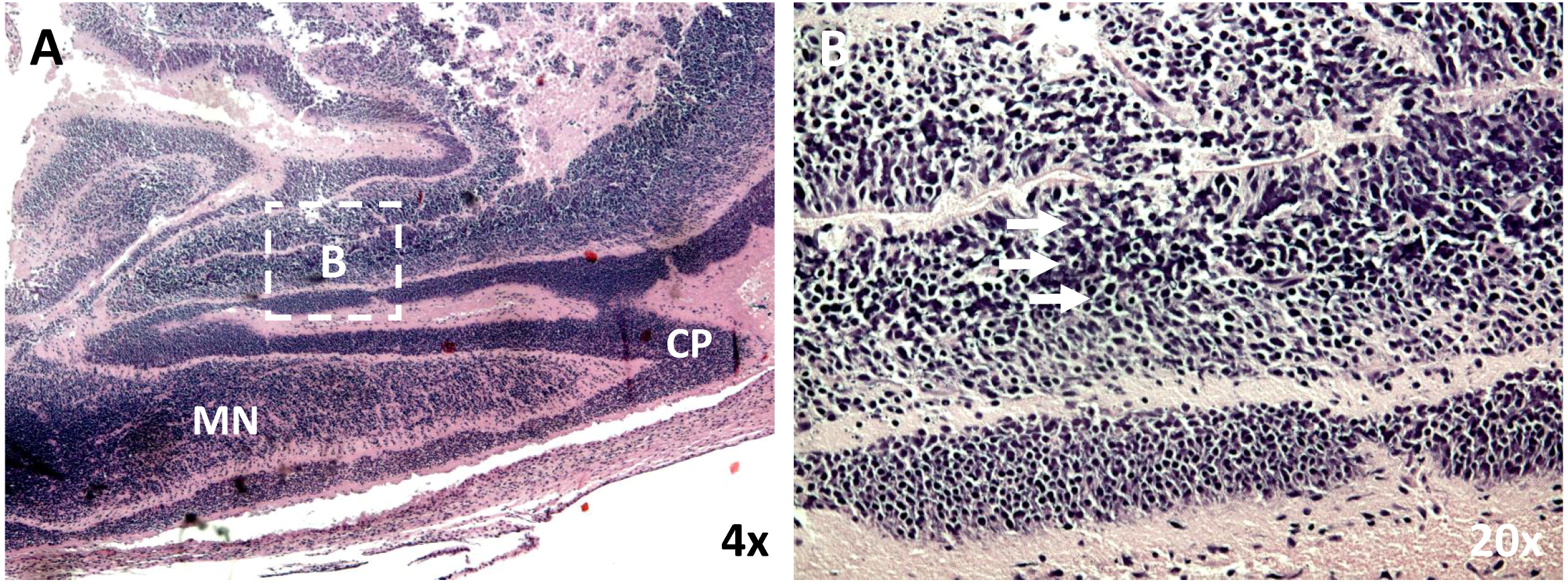
Maternal infection during placentation renders latter fetal cortical brain infection with disorganized neuronal migration. (A) Low power sagittal view of the posterior developing cranium and brain of the dam 1 fetus localizing the cortical plate (CP) and the migrating neurons (MN). (B) Higher power view of the migrating layer showing significantly disorganized migrating neurons, (arrows). The boxed area in A localizes the higher power field B.

### ZIKV genome sequencing from fetus and placenta

ZIKV genome sequencing from tissues and body fluids, done in parallel from the original inoculum, was performed to identify mutations arising after infection. In the infected placenta and singleton aborted fetus from dam 2, a single *de novo* coding mutation was identified in the matrix protein resulting in an amino acid change from phenylalanine to leucine (F252L) (**Supplementary Table S1**). Notably, this identical mutation had been previously reported in a ZIKV genome sequenced from a human fetus with microcephaly^25^, despite being absent in 500 other reference sequences deposited in GenBank (p=0.0135 for association with microcephaly by two-tailed Fisher’s Exact Test). The original ZIKV inoculum as well as placenta and fetus from dam 1 retained the original phenylalanine residue at that position.

### Whole blood maternal transcriptome profiling

Whole blood from both the asymptomatic ZIKV-infected dams collected at post-infection days 2 (for only 1 of 2 dams), 7, and 30 were collected for transcriptome profiling by RNA-seq, and compared to whole blood samples drawn from the same animals prior to ZIKV inoculation. Each sample was sequenced in duplicate to increase statistical power. The average sequencing depth was 23.9M reads per sample (±6.7M reads) (**Supplementary Fig. S1**), and STAR/Cufflinks detected an average of 55.1% (±9.8%) of all Ensembl isoforms in each sample.

We analyzed the RNA-seq data for differentially expressed genes (DEGs) at 2, 7, and 30 days post-infection relative to the uninfected baseline. Distinct from our previously published in male marmosets, in which maximal differential expression was observed at day 7 post-ZIKV infection^21^, differential gene expression in pregnant marmoset dams peaked by day 2 post-infection, returning to near baseline by day 30 (**Supplementary Table S2**). Canonical pathway analysis showed that the interferon signaling pathway was the top pathway significantly up-regulated at both days 2 and 7 post-infection (**Supplementary Fig. S2A**). Most of the differentially up-regulated genes corresponded to the type I interferon pathway, although both type I and type II pathways were predicted to be activated (**Supplementary Fig. S3**). Cytokine profiling assays showed increases in activity of a number of proinflammatory cytokines (**Supplementary Fig. S2B**). After subtracting DEGs found in common with ZIKV-infected male marmosets^21^, we used gene ontology analysis with the PANTHER database^26^ to evaluate the 29 remaining DEGs at day 7 for those that may be playing a role in the fetal loss and neuropathology (**Supplementary Table S3**). Two such candidate genes among the 29 were identified: TRPV6, a member of the TRP membrane family that plays a role in maternal-fetal Ca^2+^ transport, and LYPD6, a regulator of Wnt/β-catenin signaling involved in embryogenesis, tissue homeostasis, and regeneration.

## DISCUSSION

Rodents, including laboratory mice, are not a natural host of ZIKV infection because their IFN-driven immune responses can inhibit ZIKV replication^7^,^53,54^. As a result, murine models of ZIKV infection require IFN manipulation and are thus inherently limited in their potential for both translational relevance and mechanisms of congenital pathogenesis^7,39,53,54^. Here we show that ZIKV inoculation of pregnant marmosets causes a maternal asymptomatic infection and seroconversion similar to that seen in humans. This maternal infection is measured as prolonged viremia and viruria, and is accompanied by active viral replication in the placenta, with resultant fetal infection, a predominant neuropathology of the developing fetal brain and eye, and early demise. Although atypical for non-pregnant human cases, persistence of virus in serum and urine (i.e. viremia and viruria) beyond 14 days and as long as 8 months has been reported in ZIKV-infected pregnant women^25,27^, and is felt to be due to permissive replication in the placenta and/or fetus functioning as a viral reservoir^3,50^. In both inoculated dams, spontaneous abortion was observed 16-18 days post infection, with extensive productive viral infection in the placenta and fetal tissues documented by qRT-PCR, immunohistochemistry, and ISH for (+) and (-) ssRNA strands. We also visualized direct infection of neural progenitor cells associated with disorganized migration. The most straightforward explanation for the measured 50% expected biparietal diameter of the fetuses from dam 1 is cranial collapse following intrauterine demise; however, we also cannot rule out this finding as an early predictor of microcephaly since it is temporally and spatially accompanied by aberrant migration patterns which are hallmarks of secondary microcephaly. In 1 of the 2 inoculations, a spontaneous *de novo* mutation, F252L, shared in common with a ZIKV genome from a human case of microcephaly, was identified. Finally, maternal interferon-associated host responses and increased activity of proinflammatory cytokines were demonstrated at early as day 2 post-inoculation. Collectively, these findings reveal several intriguing and novel mechanistic insights pertaining to the potential roles of placental replication, mutation-driven viral adaptation, and maternal responses in ZIKV-mediated fetal pathogenesis.

The first trimester of pregnancy in both humans and marmosets is critical for appropriate development of the major organ systems, especially the brain and its associated neuroprogenitor cells. Gestational ages of 79-83 days (Dam 1) and 68-72 days (Dam 2) at the time of inoculation correspond to approximately 6.6 – 8.6 weeks in humans (**Fig. 2**), and is a time of crucial placental development in the marmoset. Indeed, the marmoset spends a third of its entire 143-day gestation invested in placentation. When compared to the human, which completes organogenesis by day 56 of a 266-day gestation, the marmoset will not complete organogenesis until day 80 (**Fig. 2**). Spontaneous abortion rates in the absence of congenital viral infections at this stage are hard to document, given the small sizes of the placental and fetal material being expelled; however, the available data indicate that two abortions in this gestational age range is an unexpected, spontaneous finding. For 2016-17, the SNPRC non-infected marmoset colony documented two abortions with placental/fetal material indicating this gestational age range out of 103 pregnancies (1.9%). The most comprehensive data documenting loss rates at this stage of marmoset pregnancy come from hormonal verification of pregnancy and pregnancy loss in a large (n=596) number of marmoset pregnancies in a research colony in Berlin^28^. The investigators documented a pregnancy loss rate during days 80-142 of 4.4%; our finding of 2 of 2 ZIKV-infected pregnancies resulting in spontaneous abortion is thus statistically significant (p=0.011 by Fisher’s Exact Test). In comparison, the overall loss rate in experimentally ZIKV-infected macaques at a comparable stage of embryonic development is 38%, as compared to expected non-infected loss rates of approximately 4-10% (Dudley, *et al.*, under review). In humans, documented rates of pregnancy loss due to ZIKV across all 3 stages at time of infection are estimated to be ~3%, with 8% of completed pregnancies following infection during the first trimester resulting in ZIKV-associated birth defects^4^. However, due to inherent limitations in the studies of miscarriages, early and mid-gestation ZIKV-induced pregnancy loss has been likely underreported and largely limited to case reports and limited retrospective series^36,52^. We speculate that our observation of uniform pregnancy loss in our marmoset model is a highly significant novel finding of potential translational significance and worthy of prioritized further human studies and re-analysis of existing epidemiologic data^41^.

Only rodents rendered deficient of type I interferon through experimental manipulations are highly susceptible to maternal-fetal transmission and resultant congenital disease^53,54^. It is thus of likely importance to note that in ZIKV-infected pregnant marmoset dams, maximal RNA-seq differential expression between the infected and uninfected state and significant up-regulation of interferon-associated genes in peripheral blood were observed at day 2. This early IFN systemic burst in pregnant dams is in contrast to day 9 in a previous study of ZIKV-infected male marmosets^21^. This may reflect the enhanced susceptibility of pregnant versus non-pregnant individuals to ZIKV infection, driving a more rapid host response and resulting in higher viral loads as well as persistence in body fluids and tissues^25,27^. In addition, it may provide crucial insight into the role of the placenta as both reservoir and conduit. We found early (day 2) increases in the concentrations of several proinflammatory molecules, including IFN-γ, which drives the type II interferon pathway. Interestingly, an antiviral response characterized by increased interferon and proinflammatory cytokine activity has been previously reported in association with ZIKV infection of primary human placental macrophages (Hofbauer) cells and cytotrophoblasts ^29^. Here we observed extensive ZIKV infection of the placenta and fetus in the absence of overt inflammation and despite induction of a robust host antiviral response. This is consistent with ours and others recently described human placental pathogenesis^35–38,55–57^.

Among the 31 candidate DEGs at day 7 post-infection, the decreased expression of TRPV6 and LYPD6 relative to baseline is especially intriguing. TRPV6 is highly expressed in placenta^30^, and has been shown to be critical for maternal-fetal Ca^2+^ transport and fetal Ca^2+^ homeostasis in an aromatase knockout pregnant mouse model^31^. In this mouse model, decreased expression of TRPV6 was associated with estrogen deficiency, which increases risk of miscarriage^32^. Like TRPV6, LYP6 is highly expressed in placenta^33^, and was critical for normal neuroectodermal development by way of Wnt/β-catenin signaling in zebrafish embryos^34^. Further studies are needed to determine whether viral suppression of these highly expressed maternal placental genes plays a role in the spontaneous abortions and disrupted fetal neurodevelopment seen here in association with first-trimester ZIKV infection.

An important point of consideration for applying our current findings to human congenital disease is an appreciation of the timeline of embryonic and placental development in marmoset monkeys versus humans. Marmosets have a delayed period of embryogenesis relative to humans (**Fig. 2**), resulting in the marmoset undergoing critical stages of neural development in association with a relatively far more mature placenta (both structurally and in terms of duration of the placenta as a functioning organ). Our findings further suggest that marmosets may be particularly sensitive to fetal Zika infection with respect to fetal loss, perhaps related to the initial infection of a relatively mature placenta when considered against their human counterparts (**Fig. 2**).

Our findings herein summarily show that the marmoset model faithfully recapitulates key characteristics of CZS in humans that are not observed in other rodent nor NHP models. Notably, ZIKV-infected rhesus macaques exhibited no fetal CNS abnormalities with maternal infection^15^. Pigtail macaques exhibited neural injury with severe hypoplasia and asymmetry within the occipital-parietal lobes^11^, but did not manifest the diffuse effects on neuroprogenitor cells observed in both the marmoset and human cases. In contrast, neuronal disorganization in ZIKV-infected pregnant marmosets was observed during early gestation at the posterior and crown in what will later comprise the occipital–parietal lobes (**Fig. 8**). We speculate that the capacity for the marmoset to better recapitulate human infection is related to ongoing placental development and trophoblast renewal, whereby the virus is permissive to replication in both placental trophoblasts^35^ and placental macrophages (Hofbauer cells)^36^ without evidence of inflammation^35–38,55–57^. As is likely the case in humans, the placenta may be viewed as an effective “shuttle” to the fetus, functioning first as a reservoir and later as a source of fetal infection of susceptible cell types and tissues, including neuroprogenitor cells^37^. Nevertheless, in fetuses from both dam 1 and dam 2, the virus appears to have caused injury to the neocortex documented by histopathological examination and neuronal disorganization in the absence of localized inflammation, as also seen in mouse models^7^ and human cases^38^.

For both ZIKV-infected marmosets and humans, fetal infection appears to be established in the absence of localized fetal and placental inflammation^35–38,55–57^. However, there is clear evidence of injury to the fetal neocortex as visualized by histopathological examination showing neuronal organization, mimicking findings in mice models of ZIKV vertical transmission as well as human cases^7,38^. This notion of the significance of placental mechanics and placental cell infection in the pathogenesis in CZS has been previously brought forward by a number of investigators^29,35,37,39,40,55–57^. It has long been supposed that shuttling of bloodborne ZIKV to the developing human fetus requires establishment of maternal blood in the intervillous space, which occurs approximately at 6-8 weeks post-conception (mean of roughly 42 days) (**Fig. 2**), or 8-10 weeks of pregnancy, corresponding to the late first trimester. However, epidemiologic observations reveal that both first and second trimester are highly vulnerable windows of CZS susceptibility^4,41^. It has thus been unclear whether CZS susceptibility in the first trimester may relate to leaks in placental architecture or shifting antiviral capacity of the maturing trophoblast^39^.

Our results here document a definitive role for early infection of a maturing placenta in the response to ZIKV infection during the critical period of fetal neurodevelopment (**Fig. 2**). We observed (1) active and robust ZIKV replication in the absence of local inflammation, (2) a maternal host response dominated by interferon type I and II-associated genes and proinflammatory cytokines, and (3) evidence of a functional placenta (normal amniotic fluid volume) and viable fetus despite viral replication and disruption of neuronal migration in developing brain and eye demise. Taken together, these findings suggest that contemporary ZIKV strains are capable of both replicating in the placenta and migrating to the developing fetus despite robust antiviral maternal responses. Our findings are consistent with previous work showing that multiple placental cell types, including macrophages, cytotrophoblasts, and primary human trophoblasts, are permissive to ZIKV infection ^29,35–38,55–57^. Thus, our studies in both humans and marmoset reveal that the placenta likely serves both as a conduit for fetal infection and a reservoir for ongoing replication^29,35^.

Contemporary ZIKV strains may have evolved to acquire the capacity for placental replication and fetal neuropathogenesis^29,35,39^. Notably, viral genome sequencing showed both viruses retained the S139N mutation in the pre-membane (prM) protein (associated with increased ZIKV infectivity of neural progenitor cells and more severe microcephaly in a mouse model^42^). Of note, the African strains used in the work of Bayer et al^55^, where they failed to see robust ZIKV replication in primary human trophoblasts, did not demonstrate the S139N mutation. The virus from dam 2 had a further *de novo* F252L coding mutation shared in only 1 of 6 available microcephalic ZIKV genomes in humans^25^. The serendipitous finding of a F252L coding mutation is unlikely to be coincidental, as the mutation was not seen in any of the other 603 ZIKV genomes sequenced to date not associated with microcephaly (p=0.0135). Further investigation of the potential functional significance of the F252L mutation is warranted.

We are enthused by both the potential relevance and significance of our findings employing pregnant marmosets. First, they show that the marmoset faithfully recapitulates all crucial aspects of human CZS and does not require the exogenous manipulation of interferons, necessary in murine models^7,39,53,54^. Given the capacity for the marmoset to complete multifetal gestations in 143 days and undergo repeat pregnancy within 10 days, this is an efficient model whose reproductive capacity over short time intervals rivals most rodent models. Second, these studies underscore crucial mechanistic insights pertaining to the role of the placenta in modulating (or failing to modulate) congenital infection and maternal-to-fetal transmission. Understanding the role of placental permissiveness in this process is of utmost crucial importance and reinforces the emerging notion of the placenta not as a *de facto* barrier, but as an active conduit for maternal and fetal communication during gestation. Thus, the identification of drugs not only to treat or prevent ZIKV infection of the fetus but also to prevent placental infection and viral transport may be critical to prevent the irreversible neuronal damage associated with CZS. Third and of potential crucial clinical importance, our findings demonstrate a high rate of pregnancy loss with maternal ZIKV infection and placental and fetal transmission. These observations should prompt reexamination of human epidemiologic data and provide additional endpoints in prospective studies as ZIKV continues to emerge in newly endemic communities in Texas, Florida, and other at-risk regions of the Americas.

## METHODS

### Marmoset inoculation and sample collection

Animal studies were conducted at the Southwest National Primate Research Center (SNPRC), an AAALAC-accredited program. All studies were reviewed and approved by the local Institutional Animal Care and Use Committee (IACUC) and Biohazard Committee.

Two pregnant dams (4.2 and 3.8 years of age; 474 and 422 grams body weight) were inoculated intramuscularly with 0.25 mL containing with 2.5x10^5^ plaque forming units (PFU) of the Brazil ZIKV strain SPH2015 diluted in saline (GenBank accession number KU321639)^43^, followed by a second identical inoculation 4 days later. The gestational age of the embryos at the time of inoculation was determined by crown-rump length, which we have previously demonstrated to provide a reliable gestational age to within +/- 3 days^44^. Embryonic growth was tracked by ultrasonography performed on unsedated animals. For dam 1, ultrasound exams took place on days 80, 88, and 93, confirming two live embryos with biparietal diameters as expected for those ages. Ultrasounds on dam 2 were on days 78 and 82, confirming one live embryo. In each case, the number of embryos was consistent from pre-dosing screening through the infection period.

Following ZIKV infection, pregnant dams were monitored daily for symptoms. Both females gained weight over the period from inoculation to abortion (0.4 grams −1.5 grams per day), within or exceeding the published range of weight gain for pregnant dams at mid-term (0.39 – 0.71 grams/day)^45^. Urine, blood, and saliva were sampled and serially monitored for ZIKV RNA by qPCR testing per a pre-established schedule (**Fig. 1A**). For the 2 female marmosets, blood samples to obtain serum were collected via venipuncture on days −5, 2, 7 and 30; voided urine was collected on days −5, 2, 7, 9, 10, 14, 21, and 28; saliva was collected on days −5, 2, 7, 14, 21, 28 and 30 by allowing the subjects to chew on a sterile cotton swab. Whole blood samples for transcriptome analysis were collected for dam 1 on days 7 and 30 and for dam 2 on days −5, 2, 7, and 30 in tubes containing RNA stabilization media (Biomatrica, Inc.). At day 30, both female marmosets were euthanized and their necropsy tissues examined for persistent ZIKV infection (**Fig. 1A**).

### Measurement of ZIKV RNA loads by quantitative RT-PCR

The course of infection in inoculated marmosets was monitored by determination of ZIKV RNA loads (expressed as RNA copies / mL) in infected body fluids (serum, urine, and saliva) and placental and fetal tissues. ZIKV RNA loads were calculated by generation of a standard curve, followed by quantitative RT-PCR testing for 45 cycles using published PCR conditions and primers targeting the envelope gene (ZIKV-1086/ZIKV-1162)^5^. By standard curve analysis, the estimated limit of detection for the qRT-PCR assay was ~15 RNA copies per mL.

### ZIKV serological analysis by antibody neutralization

Plaque-reduction neutralization assay (PRNT) on longitudinally collected marmoset sera was performed by the California Department of Public Health, using a protocol adapted from that used by the US Centers for Disease Control and Prevention (CDC) for confirmatory ZIKV testing in patients. Briefly, 100 plaque forming units (PFU) of ZIKV (Brazilian SPH2015 strain) were mixed with equal volumes of serial 2-fold dilutions of inactivated marmoset sera and incubated for 1 hr at 36°C, followed by inoculation and adsorbing to a monolayer culture of Vero cells for 1 hr at 36°C. After addition of 3 mL of 0.6% agar in Eagle’s Minimal Essential Medium (MEM), plates were placed in a 36°C, 5% CO_2_ incubator for 3 days, followed by addition of 3 mL of 1% agar and 0.004% neutral red in Eagle’s MEM and another 1-2 days of incubation until plaques were formed. An 80% or more reduction of the number of plaques compared to positive control wells inoculated with virus-diluent mixtures was considered neutralization, with serum titers reported as the highest dilution exhibiting 80%or more reduction.

### Deep sequencing analysis of ZIKV recovered from marmoset organs and body fluids

Nucleic acid was extracted from 50 μg of placenta or fetus lysate, 50 μL of body fluid sample (serum, urine, or saliva), or 100 μL of viral culture inoculum using the automated Qiagen EZ1 instrument (Qiagen, Valencia, CA), followed by treatment with Turbo DNase (Thermo-Fisher, Carlsbad, CA). cDNA was obtained by reverse transcription using a 1:5 mix of random hexamer and short spiked primers targeting ZIKV genomes (ref). Next-generation sequencing (NGS) libraries were constructed using the Nextera XT kit (Illumina, San Diego, CA), followed by single-end, 150 base pair (bp) sequencing on an Illumina MiSeq or HiSeq2500 instrument. Raw reads were preprocessed by adapter trimming and removal of low-quality, low-complexity sequences as previously described^46^. Data was scanned for ZIKV reads using an NCBI BLASTn database constructed from all known ZIKV sequences as of October 2017, using an E-value significance threshold of 1x10^−5^. Consensus ZIKV genomes were assembled using Geneious (v10.2.2) by mapping preprocessed ZIKV reads to reference genome KU321639.

### Fetal and placental histology and high-resolution imaging

A full sagittal section was mounted of one dichorionic fetus from dam 1, and frontal and coronal sections of the singleton dam 2 pregnancy were formalin-fixed and paraffin-embedded. Sections were H&E stained to assess for localized inflammation, as well as to visualize and identify tissues within the fields. Serial sections of all marmoset tissues were further probed for ZIKV viral genome using RNAscope ZIKV probes (Advanced Cell Diagnostic, Inc.) for both + (stable) and – (active replicating) RNA strands to determine subcellular localization of viral establishment and active replication. Probing utilized established methodologies^36^. Briefly, 4 µM sections were deparaffinized and subjected to antigen retrieval as per manufacturer’s instructions (RNAscope 2.5 HD Detection Kit, ACD). A previously designed probe set of complementary oligos against either strand, covalently bound to adapter sequences was then applied^36^. The ZIKV sequence specific probe sets consisted of 87 probe pairs requiring hybridization of adjacent matching genome sites for adapter probe hybridization and amplification. DAB signal was generated after a series of amplification probes as per manufacturer’s instructions. Placenta were also stained with 4G2 antibody against flavivirus protein E (Millipore) as previously described^35^. Vero cells (Green monkey kidney epithelial) infected with contemporaneous ZIKV were used as a positive control. Cells were infected with first passage ZIKV HN16 (GenBank: KX928077.1) isolated from traveler returning to the USA from Honduras. Following 6 days of *in vitro* infection and replication, scraped cells were formalin fixed in a syringe, then paraffin embedded. Bright field slides were examined under low and medium power using a Nikon Eclipse 90i microscope (Nikon Instruments, Melville, NY, USA). Minor adjustment for contrast and background levels were made using NIS Elements Viewer 4.20 (Nikon) and Adobe Photoshop CC.

### Multiplex cytokine analysis of plasma

Plasma samples were analyzed for marmoset cytokines and chemokines by Luminex using established protocols for New World primates (Giavedoni, LD. Journal of Immunological Methods 301, 89–101, 2005). Evaluation of analytes GRO-α (CSCL1), interferon alpha (IFN-α), IFN-γ, interleukin-1 beta (IL-1β), IL-22, monocyte chemoattractant protein 1 (MCP-1, CCL2), macrophage migration inhibitory factor (MIF), monokine induced by gamma interferon (MIG, CXCL9), macrophage inflammatory protein 1-alpha (MIP-1α, CCL3), MIP-1β (CCL4), regulated on activation, normal T cell expressed and secreted (RANTES, CCL5), tumor necrosis factor-alpha (TNF-α), soluble CD40 ligand (sCD40L), soluble intercellular adhesion molecule 1 (sICAM-1), and vascular endothelial growth factor A (VEGF-A) were included in this assay. Only analytes quantifiable above the limit of detection are presented.

### Transcriptome analysis

Prior to inoculation, whole blood was drawn from the two age-/sex-matched healthy pregnant dams in the ZIKV study to use as internal controls in the transcriptome analysis. All marmosets studied here were from a single colony, thus increasing genetic similarities and decreasing environmental bias. 400 uL microliters of blood were drawn directly into RNAgard tubes (Biomatrica) for immediate RNA stabilization of intracellular RNA at time of collection. Total RNA was extracted using the Biomatrica Blood RNA Purification Kit (Biomatrica). The Ovation Human Blood RNA-Seq Kit (Nugen, San Carlos, CA) was used to generate RNA-seq libraries from 100 ng of input per sample (as measured using the Invitrogen Qubit RNA HS Assay Kit) according to the manufacturer’s protocol. Libraries were sequenced in duplicate as 100 bp paired-end runs on an Illumina HiSeq 2500 instrument.

Paired-end reads were mapped to the marmoset genome (Callithrix jacchus Ensembl version 3.2.1), using STAR 2.5^47^, and gene and transcript normalized counts were calculated by HTSeq version 0.6.0^48^. Differential expression of genes was calculated using linear modeling using the Bioconductor EdgeR software package version 3.12.2^49^, as implemented in the R programming language. Genes were considered to be differentially expressed when their fold change was > ± 2, *p*-value < 0.05, and adjusted *p*-value (or false discovery rate, FDR) < 0.1%. Pathway and network analyses of the transcriptome data were performed using Ingenuity Pathway Analysis (IPA) software (Qiagen). Marmoset transcriptome data has been submitted to the public National Center for Biotechnology Information (NCBI) Gene Expression Omnibus (GEO) repository (accession numbers pending).

### Statistical Tests

All p-values reported in this study are calculated using two-tailed Fisher’s Exact Test.

## CONFLICTS OF INTEREST

The authors disclose no conflicts of interest.

## ACKNOWLEDGEMENTS

We would like to acknowledge Lark Coffey from UC Davis and Marion Lanteri from Blood Systems Research Institute for kindly sharing their P3 stock of ZIKV Brazil strain SPH2015. We also thank Julien Thézé at Oxford University for assistance in reviewing available ZIKV reference genomes in GenBank. We thank Anthony Chiu for line art drawing of the embryo images shown in Figure 8. The animal work in this study was supported by the Southwest National Primate Research Center (P51-OD11133) and Texas Biomedical Research Institute. This study was supported in part by NIH grants R01-HL105704 and R21-AI129455 to CYC and R24-DK090964, R01-HD091731, and R01 DK089201 to KMA. The funders had no role in study design, data collection and analysis, decision to publish, or preparation of the manuscript.

## AUTHOR CONTRIBUTIONS

MS, CS-SM, SDT, KMA, CYC, and JLP conceived and designed the experiments. MS, MF, CS-SM, TL, VLH, LMP, LG, DL, MAS, KMA and MT performed the experiments. MS, CS-SM, SDT, JR, TL, LG, KMA, CYC, and JLP analyzed the data. LG, MAS, KMA, CYC, and JLP contributed reagents/materials/analysis tools. MS, CS-SM, SDT, JR, LG, KMA, CYC, and JLP wrote the paper. MS, SDT, JR, LG, KMA, and CYC prepared the figures. All authors reviewed the manuscript and agree to its contents.

